# First report of astroviruses in Tanzanian bats

**DOI:** 10.1101/2024.03.05.581362

**Authors:** Léa Joffrin, Evangelia Iliopoulou, Marta Falzon, Christopher Sabuni, Lucinda Kirkpatrick, Luc De Bruyn

**Affiliations:** Evolutionary Ecology Group, Department of Biology, University of Antwerp, Universiteitsplein 1, 2610 Antwerp, Belgium; Pest Management Institute, Sokoine University of Agriculture, Morogoro, Tanzania; Research Institute for Nature and Forest (INBO), Havenlaan 88 bus 73, 1000 Brussels, Belgium; School for Environmental and Natural Sciences, Bangor University, Bangor, LL57 2DG Wales

**Keywords:** astrovirus, bat, Tanzania, virus, phylogeny

## Abstract

Emerging and re-emerging infectious diseases have posed significant global health threats, with many attributed to zoonotic RNA viruses. These pathogens can, under some conditions, cross species barriers, facilitating transmission from animal hosts to humans. Bats, characterised by unique physiological and ecological features, and remarkable species diversity, are recognized to host numerous viruses with cross-species transmission potential. This study aimed to investigate the presence of RNA viruses from a broad diversity of Tanzanian bats while valorising archived biological samples. RNA was extracted from 125 samples (28 faeces and 97 oral swabs) of 17 bat species, followed by PCR amplification targeting five distinct viral genera (Filovirus, Coronavirus, Hantavirus, Paramyxovirus and Astrovirus). Overall, 1.6 % (3/125) of the samples from two bat species (*Scotophilus dinganii* and *Miniopterus fraterculus*) tested positive for astrovirus, with the coinfection of one bat with two AstV strains. No samples tested positive for Filovirus, Coronavirus, Hantavirus and Paramyxovirus. Phylogenetic analysis based on RNA-dependent RNA polymerase sequences revealed these sequences are respectively clustering with astroviruses detected in other bat species from the genus Scotophilus from East Asia and with astroviruses detected in *Miniopterus* bats from Africa and Asia. Altogether, these results are the first report of astroviruses in Tanzanian bats.

## Introduction

Zoonotic pathogens originating from wildlife, and especially RNA viruses, are the main cause of emerging infectious diseases (1). RNA viruses have a high mutation rate compared to other viruses, allowing for rapid adaptation to potential other hosts. Bats are the natural reservoir of many infectious agents, including RNA viruses that cause severe human diseases such Nipah virus or Hendra virus (2,3). Many of these RNA viruses are thought to be mainly transmitted via contact with bat excreta such as saliva, urine, and faeces (4).

Bat diversity in East Africa is among the highest in the world, and Tanzania harbours over a hundred bat species, with six of them endemic to the country (5). However, the surveillance of bat-borne viruses has been focused mostly on Asian bats, with twice as many viral sequences reported from Asian bats than African bats and 60% of Asian sequences isolated in China (6). Early investigations on East African bats have already identified a great diversity of RNA viruses (such as Filoviridae, Paramyxoviridae, and Coronaviridae) in bats from Kenya, Uganda, Rwanda, Mozambique, Ethiopia, West Indian Ocean islands (7–13). In addition, viruses of potential public health concern like SARS-related CoVs and henipa-related paramyxoviruses have also been described in bats from East Africa (7–10). Despite more than 650 viral sequences isolated from Tanzanian bats between 2010 and 2018, investigations mainly were focused on coronaviruses (99% of the detected sequences) and paramyxoviruses (10,14,15).

In this study, we tend to valorise archived biological samples from a broad diversity of Tanzanian bats. While these samples were initially collected for species identification and diet analysis, we screened them for five viral families and tried to assess if viral RNA could still be detected in these samples despite suboptimal storage and conservation medium.

## Material and method

Bats were captured with mist nets and harp traps from July to October 2018 in different locations in Tanzania (Morogoro, Saadani National Park (NP), Udzungwa NP and Ruaha NP) for other projects to assess Tanzanian bat diversity and their diet. Bats in the field were identified based on external morphological characteristics and based on Monadjem et al. (2011) (16). Trapping locations, sex, age, and reproductive status were recorded. Biological samples were collected for molecular bat species barcoding and diet analysis. Briefly, 3mm diameter wing punches and faeces samples were stored on silica beads. Oral swab samples were kept in RNAlater™ stabilisation solution. All samples were kept at ambient temperature for several days when sampling occurred, except for samples collected in Morogoro, where the samples could be stored in a fridge. After arrival in Belgium, the samples were stored at the EVECO laboratory at the University of Antwerp. The faecal samples and oral swabs were stored at - 20°C.

Each faecal sample was homogenised in 500 μL of PBS in a class II biosafety cabinet. Oral swabs and faecal samples were then vortexed and centrifuged at 1500g for 15min. RNA extraction was performed using the QIAamp Viral RNA Mini Kit (QIAGEN, Valencia, CA, USA). Reverse transcription was performed on 8 μL of the extracted RNA as described in (17). cDNas were tested for the presence of five distinct viral families (Filovirus, Coronavirus, Hantavirus, Paramyxovirus and Astrovirus) using pan-PCR systems previously published and targeting highly conserved regions of RNA-dependent RNA-polymerase (RdRp) genes (11,12,18–20). All PCR products of the expected size were submitted for direct Sanger sequencing (Neuromics Support Facility of the VIB-UAntwerp Center for Molecular Neurology, Antwerp, Belgium).

Genetic diversity of the sequences was explored with pairwise identity values and BLASTn on Geneious software 2022.1.1 (21). The sequences were edited and aligned with 122 partial AstV RdRp sequences representing a large diversity of hosts and geographic origins (Africa, Asia, Europe, Oceania, America) using Geneious software 2022.1.1 (21). A phylogenetic tree was obtained by maximum-likelihood using PhyML (22), with 1,000 bootstrap iterations. The best evolutionary model for the phylogenetic reconstruction selected by Model Generator v0.8565 (23) based on Akaike Information Criterion 1 (AIC1) was GTR, with a proportion of invariable sites of 0.11 and the gamma distribution parameter α = 0.83.

## Results & Discussion

In total, 125 samples from 17 different bat species were tested, with 97 oral swab samples from 17 different bat species and 28 faecal samples from 8 different bat species (**Table**). No samples tested positive for Filovirus, Coronavirus, Hantavirus and Paramyxovirus. Astroviruses (AstV) were detected in one *Scotophilus dinganii* from the Vespertillionidae family and one *Miniopterus fraterculus* from the Miniopteridae family, both trapped in Morogoro. While astroviruses also have been detected in Scotophilus bats from Asia, no astroviruses has been reported from a *Scotophilus* bat from Africa. Also, no astroviruses has been reported from any other bat family in Tanzania (6). We obtained three partial RdRp sequences (422 nt). Pairwise comparison of the three sequences revealed three unique sequences, with similarities ranging from 60.9% to 76.8%. The two sequences of *Miniopterus fraterculus* and the one from *Scotophilus dinganii* show the lowest sequence similarity (60.9% and 64.2%). The two sequences isolated from *Miniopterus fraterculus* share 76.8% nucleotide similarity. A maximum-likelihood phylogenetic reconstruction was performed using these sequences and 125 AstV RdRp partial nucleotide sequences. The 422 nt AstV sequence detected in *Scotophilus dinganii* shares 77.3% nucleotide similarity with an AstV detected in *Scotophilus kuhlii* from China (HQ613175). This sequence clustered in a monophyletic group formed by AstVs of *Scotophilus* bats from East Asia (**Figure**). The clustering of this AstV with Asian bats may come from a lack of AstV sequences from *Scotophilus* bats from Africa compared to Asia since 73% of all the sequences published have been detected in Asia and only 19% in Africa (6). We detected two AstV variants in the same samples from *Miniopterus fraterculus*. One 422 nt AstV sequence shares 83.9% nucleotide similarity with an AstV detected in *Miniopterus sororculus* from Madagascar (MZ614418) and clusters with other sequences detected in *Miniopterus* bats in Africa and Asia (**Figure**). The other one shares 82.5% nucleotide similarity with an AstV detected in *Miniopterus magnater* from Thailand (KY054113) and clusters together with other sequences detected in *Miniopterus* bats (**Figure**). Interestingly, these sequences are closer to sequences from Malagasy bats than sequences isolated in other species in continental East Africa (Kenya, Mozambique) (13,26). However, the low nucleotide similarity percentage of our sequences with reference sequences, along with the absence of robust statistical support for certain clusters in the phylogenetic tree, indicate that the small number of sequences identified in Africa may not provide sufficient data for a reliable comparison of AstV evolution and ecology when contrasted with the broader spectrum of astroviruses discovered in Asia.

**Table.** Total number of samples screened per bat species and sample type. In red, the number of samples detected positive for astrovirus.

| Bat taxa | Npos/Ntotal samples screened | Npos/N faeces tested | Npos/N oral swab tested |
| --- | --- | --- | --- |
| **Megadermatidae** | **6** | **3** | **3** |
| *Lavia frons* | 6 | 3 | 3 |
| **Miniopteridae** | **2** | **0** | **2** |
| *Miniopterus fraterculus* | 1/2 | 0 | 1/2 |
| **Molossidae** | **39** | **17** | **22** |
| *Chaerephon pumilus* | 39 | 17 | 22 |
| **Nycteridae** | **2** | **1** | **1** |
| *Nycteris hispidus* | 2 | 1 | 1 |
| **Pteropodidae** | **47** | **2** | **45** |
| *Epomophorus crypturus* | 1 | 0 | 1 |
| *Epomophorus grandis* | 1 | 0 | 1 |
| *Epomophorus labiatus* | 28 | 2 | 26 |
| *Epomophorus wahlbergi* | 10 | 0 | 10 |
| *Epomophorus grandis* | 1 | 0 | 1 |
| *Epomophorus* sp. | 4 | 0 | 4 |
| *Rousettus aegyptiacus* | 2 | 0 | 2 |
| **Rhinolophidae** | **4** | **1** | **3** |
| *Rhinolophus clivosus* | 3 | 1 | 2 |
| *Rhinolophus eloquens* | 1 | 0 | 1 |
| **Rhynonycteridae** | **5** | **2** | **3** |
| *Triaenops persicus* | 5 | 2 | 3 |
| **Vespertilionidae** | **20** | **2** | **18** |
| *Glauconycteris argentata* | 2 | 1 | 1 |
| *Neoromicia nana* | 14 | 1 | 13 |
| *Neoromicia rendalli* | 1 | 0 | 1 |
| *Scotophilus dinganii* | 1/3 | 0 | 1/3 |
| **TOTAL** | **2/125** | **28** | **2/97** |

**Figure.**
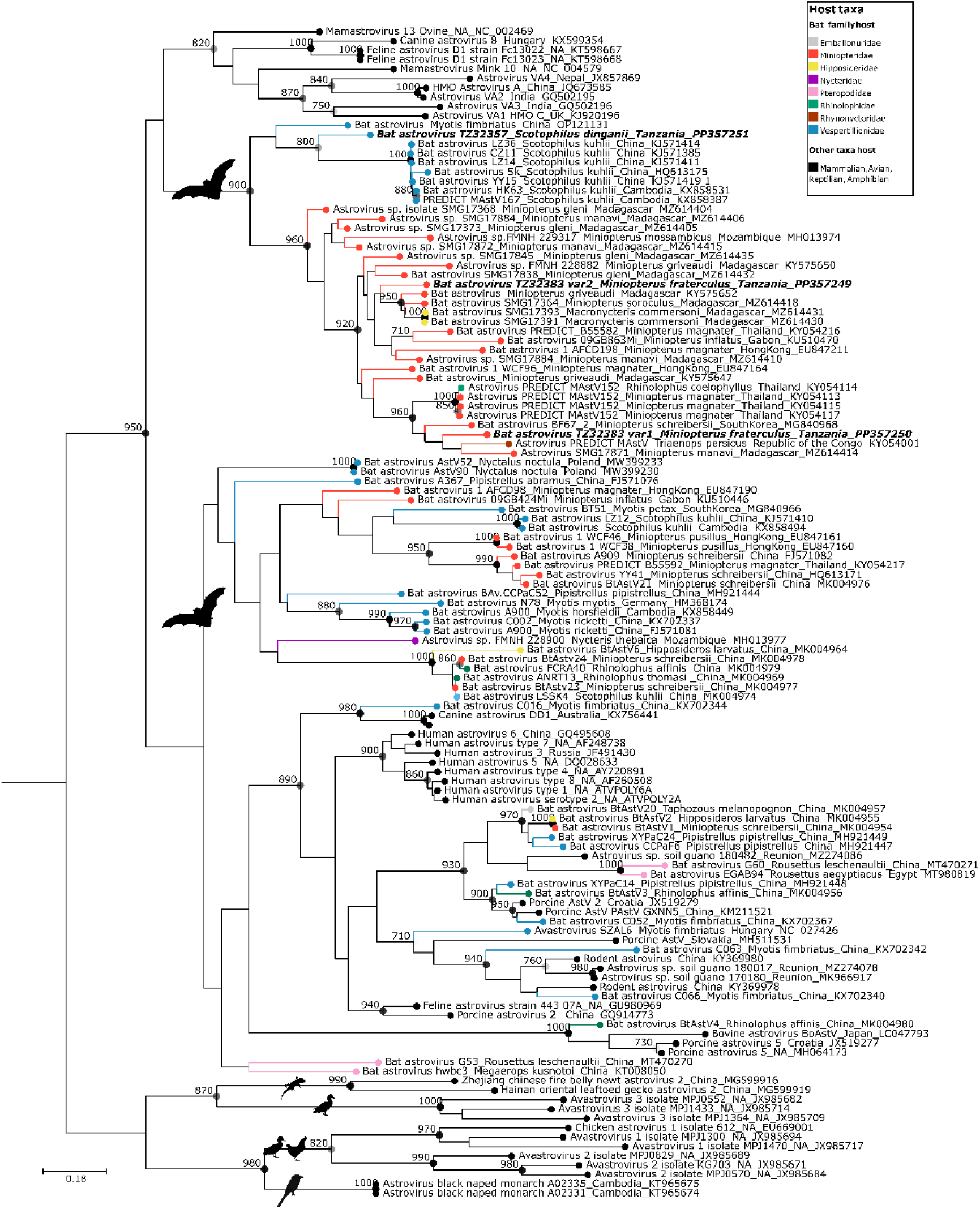
Maximum likelihood consensus tree derived from 125 Astrovirus (AstV) RNA-dependent RNA-polymerase partial nucleotide sequences (394 nt). Black dots indicate nodes with bootstrap values higher or equal to 700. Bold font: sequences generated in this study. Scale bar: mean number of nucleotide substitutions per site.

AstVs belong to the Astroviridae family and are RNA viruses infecting a wide range of mammalian and avian species (27), including numerous bat species in Asia, Europe, and Africa (13,28–36). In humans, they are transmitted via the faecal-oral route and are responsible for a significant proportion of non-bacterial enteric diseases, especially in young children (37,38). The occasional detection of avian and canine astroviruses in humans suggests a potential underestimation of interspecies transmission events (39). Studies of AstV evolution have revealed minimal host restriction among bats, suggesting host-switching events as an important driver in their evolution (11,13,40). Thus, the risk of emergence and epidemics caused by AstVs of zoonotic origin requires more knowledge about the eco-epidemiology and evolution of these viruses within reservoirs (27,38). In addition, the frequent co-infection of AstVs with other viruses of sanitary interest, such as coronaviruses, has been reported (24,26). A critical knowledge gap exists regarding the circulation dynamics of AstVs not only within bat populations but also at a broader scale within animal communities (41). Several studies have provided evidence of temporal variations in the detection rate of AstVs in bats, suggesting a potential seasonal dynamic (24,42), while others have not corroborated this observation (43). For instance, an investigation conducted in Malaysian Borneo documented a higher frequency of Astrovirus detection in bats during the rainy season compared to the dry season (28). Conversely, a meticulous two-year longitudinal study tracking astroviruses within a single-species colony on Reunion Island revealed no statistically significant variations across various sampling collection periods (35). These findings collectively emphasise the complexity of astrovirus epidemiology in bat populations and underscore the need for further research to elucidate the factors driving these observed temporal dynamics.

Overall, two oral swab samples tested positive for astrovirus (2/125). This detection rate was lower than those reported in other studies using the same PCR assay (8,11,13,24,25). However, since the mean detection rate of these viral families in bats can vary significantly depending on the region, bat species, sampling methods, and the sensitivity of the diagnostic techniques used, our result may also reflect the small sample size available for the study. Moreover, the samples we screened were initially collected for species identification and diet analysis studies. Thus, the storage conditions were not optimum for properly conserving viral RNA. Faecal samples were stored and dried on silica beads, while oral swabs were kept in RNA later, an appropriate medium for viral RNA conservation. However, the samples were not immediately frozen and were kept in suboptimal temperature conditions for several days. We thus consider it very likely that conservation issued between the collection and the final storage of samples in freezers had caused severe damage to the samples, with potential partial or total degradation of the RNA. These results stress the critical importance of utilizing appropriate conservation mediums for sample collection. Additionally, it highlights the potential biases that may be introduced when relying on opportunistic samples collected within the framework of other research projects. Nevertheless, these results indicate that further investigation of astroviridae dynamics within bat populations in Tanzania might offer valuable research opportunities to better understand astrovirus circulation in wild animal populations.

## Conclusions

The examination of these intricate viral interactions within bat populations and broader animal communities can provide valuable clues regarding the selective pressures shaping these viruses, their adaptation to different host species, and the potential risks of spillover events. Such knowledge is crucial for mitigating public health threats posed by AstVs and for implementing effective strategies to prevent future cross-species transmissions. Furthermore, it underscores the importance of continued surveillance and research efforts to unravel the complex dynamics of bat-borne viruses in the context of global health security.

## Data availability

All the data generated during the current study are included in the manuscript. DNA sequences: GenBank accessions PP357249 to PP357251.

## Ethics statement

Bat sampling was conducted under ethical permission from the Sokoine University of Agriculture. Animals were handled to minimize any potential stress or harm to the animals and were realised after. The sampling of faeces and oral swabs from bats was conducted in accordance with ethical considerations and best practices for animal welfare.

## Author contributions

Léa Joffrin: Conceptualisation, Methodology, Validation, Formal analysis, Investigation, Data curation, Writing-original draft preparation, Writing-review and editing, Visualisation, Supervision, Project administration, Funding acquisition. Evangelia Iliopoulou: Formal analysis, Investigation, Writing-review and editing. Marta Falzon: Investigation. Christopher Sabuni: Resources. Lucinda Kirkpatrick: Writing-review and editing, Funding acquisition. Luc De Bruyn: Conceptualisation, Writing-review and editing, Supervision.

## Conflict of interest

The authors declare no conflict of interest. The funders had no role in the design of the study; in the collection, analyses, or interpretation of data; in the writing of the manuscript; or in the decision to publish the results.

## Acknowledgements

We are grateful to Herwig Leirs, Laura Cuypers, the staff (in particular Shabani Lutea, Geofrey Sabuni, Baraka E. Mwamundela and Joshua Jakoniko) from the Pest Management Institute (Sokoine University of Agriculture, Morogoro, Tanzania) and the students of the Tropical Field Course Tanzania for their assistance during the fieldwork.

## Funding

LJ is a postdoctoral fellow of the Research Foundation–Flanders (FWO) [Grant # 1271922N]. MF was supported by a “Master Mind Scholarship” of the Flemish Ministry of Education and Training. LK was a postdoctoral fellow of the Research Foundation–Flanders (FWO) [Grant #1220820N]. Tropical Field Course Tanzania was partially funded by the University of Antwerp.

## Notes

### Competing Interest Statement

The authors have declared no competing interest.

### Summary of Updates

Affiliation address for the Research Institute for Nature and Forest (INBO)

